# INFERNO - INFERring the molecular mechanisms of NOncoding genetic variants

**DOI:** 10.1101/211599

**Authors:** Alexandre Amlie-Wolf, Mitchell Tang, Elisabeth E. Mlynarski, Pavel P. Kuksa, Otto Valladares, Zivadin Katanic, Debby Tsuang, Christopher D. Brown, Gerard D. Schellenberg, Li-San Wang

## Abstract

The majority of variants identified by genome-wide association studies (GWAS) reside in the noncoding genome, where they affect regulatory elements including transcriptional enhancers. We propose INFERNO (INFERring the molecular mechanisms of NOncoding genetic variants), a novel method which integrates hundreds of diverse functional genomics data sources with GWAS summary statistics to identify putatively causal noncoding variants underlying association signals. INFERNO comprehensively infers the relevant tissue contexts, target genes, and downstream biological processes affected by causal variants. We apply INFERNO to schizophrenia GWAS data, recapitulating known schizophrenia-associated genes including *CACNA1C* and discovering novel signals related to transmembrane cellular processes.

## Background

Genome-wide association studies (GWAS) have successfully identified over 50,000 genetic variants associated with more than 2,300 human diseases and phenotypes [1, 2], but the interpretation of these associations remains limited. First, each GWAS-identified variant tags linkage disequilibrium (LD) blocks of potentially functional variants that are inherited together, and the causal variant underlying the association signal may not be a genotyped variant [3]. Second, 90% or more of GWAS variants are in the noncoding genome and do not directly affect coding sequences of messenger RNAs (mRNA) [4]; rather, they may affect regulatory elements that modulate mRNA transcription levels such as enhancers [5]. Enhancers are context-specific and annotations are incomplete, so information must be integrated across tissue contexts and data sources to identify variants affecting enhancer function [6]. Finally, to translate GWAS findings into a deeper understanding of pathology leading to the development of novel therapeutics, it is crucial to identify not only the genes targeted by affected enhancers but also the tissue or cellular context in which their expression is altered that underlies disease risk. Recent large-scale efforts focused on identifying active regulatory regions within the noncoding genome [7-9], but the field lacks a comprehensive method for identifying not only causal noncoding variants and the regulatory elements they disrupt but also the relevant tissue context, target genes, and downstream biological processes affected by these variants. Researchers have developed several tools for investigating noncoding genetic signals such as RegulomeDB, GWAVA, CADD, and GenoSkyline [10–13]. These methods are unified by their approach of generating scoring functions across the genome to identify causal variants, regulatory loci, and in some cases, the relevant tissue contexts. However, while using these types of integrative scoring functions to summarize noncoding genetic function enables the ranking and identification of individual variants, these tools do not identify both the specific affected regulatory elements and the affected target genes. Another tool is HaploReg [14], which expands GWAS tag variants into haplotype blocks and overlaps them with chromatin state annotations and eQTL results to identify specific regulatory loci and target genes. However, it offers no way to integrate the enhancer and eQTL overlap results to characterize the affected tissue contexts, and performs direct eQTL overlap, which is biased by LD structure and may yield both false positives and negatives.

In this paper, we introduce the INFERNO method (INFERring the molecular mechanisms of NOncoding genetic variants), which integrates hundreds of diverse functional genomics data sources across tissues and cell lines with GWAS summary statistics to identify sets of putatively causal noncoding variants underlying an association signal and comprehensively characterize the downstream regulatory effects of these variants. INFERNO includes a tissue classification scheme that is used to integrate information across diverse functional genomics data sources. This enables the identification of variants with concordant support from multiple data sources in a specific tissue context in a hypothesis-free manner. INFERNO also introduces a novel statistical model for quantifying the enrichment of enhancer overlaps in specific tissue categories for any GWAS data.

To identify the tissue-specific affected target genes, INFERNO integrates expression quantitative trait loci (eQTL), variants whose alleles are correlated with changes in the expression level of a target gene, from the GTEx consortium [15] with GWAS summary statistics by applying a Bayesian co-localization model [16]. This allows the method to avoid the biases of directly overlapping GWAS variants with eQTL measurements and identify eQTL signals that are strongly co-localized with association signals. Furthermore, it enables the identification of functional variants that underlie co-localized signals and also overlap functional regulatory elements in the matching tissue category. Many eQTL signals affect long noncoding RNAs (lncRNAs) which in turn can regulate protein-coding gene expression, so INFERNO identifies lncRNA target genes and downstream biological processes using GTEx RNA sequencing data [15].

To demonstrate the utility of INFERNO, we applied the method to a large schizophrenia GWAS dataset [17]. INFERNO identified significant enhancer enrichments in immune-related tissue categories, uncovered functionally supported variants underlying eQTL signals targeting known schizophrenia genes including *CACNA1C* and novel candidates related to transmembrane cellular signaling. We also identified affected lncRNAs in the brain and immune system and characterized the tissue-specific downstream effects of these lncRNAs at the level of biological pathways, identifying tissue-specific effects on several processes including *MAPK* signaling, splicing, and Herpes simplex infection. INFERNO is available as an open source software package, and users can analyze top GWAS variants using the web server at http://inferno.lisanwanglab.org/.

## Results

### Overview of INFERNO pipeline

INFERNO consists of four analysis stages (Figure 1): 1. Define exhaustive sets of all potentially causal variants underlying each top GWAS signal. 2. Characterize these variants by overlapping them with various functional genomics data sources including epigenomic states, enhancer annotations, and overlap with messenger RNA (mRNA) and repeat elements. 3. Use GWAS and eQTL data for Bayesian co-localization analysis to identify tissue-specific effects on target genes. 4. Integrate information from the previous steps using a tissue categorization framework to identify functional variants with concordant annotation support, the tissue contexts of enhancer-gene interactions, target genes with strong functional support, and the downstream biological processes affected by disruption of target genes and long noncoding RNAs.

**Figure 1:**
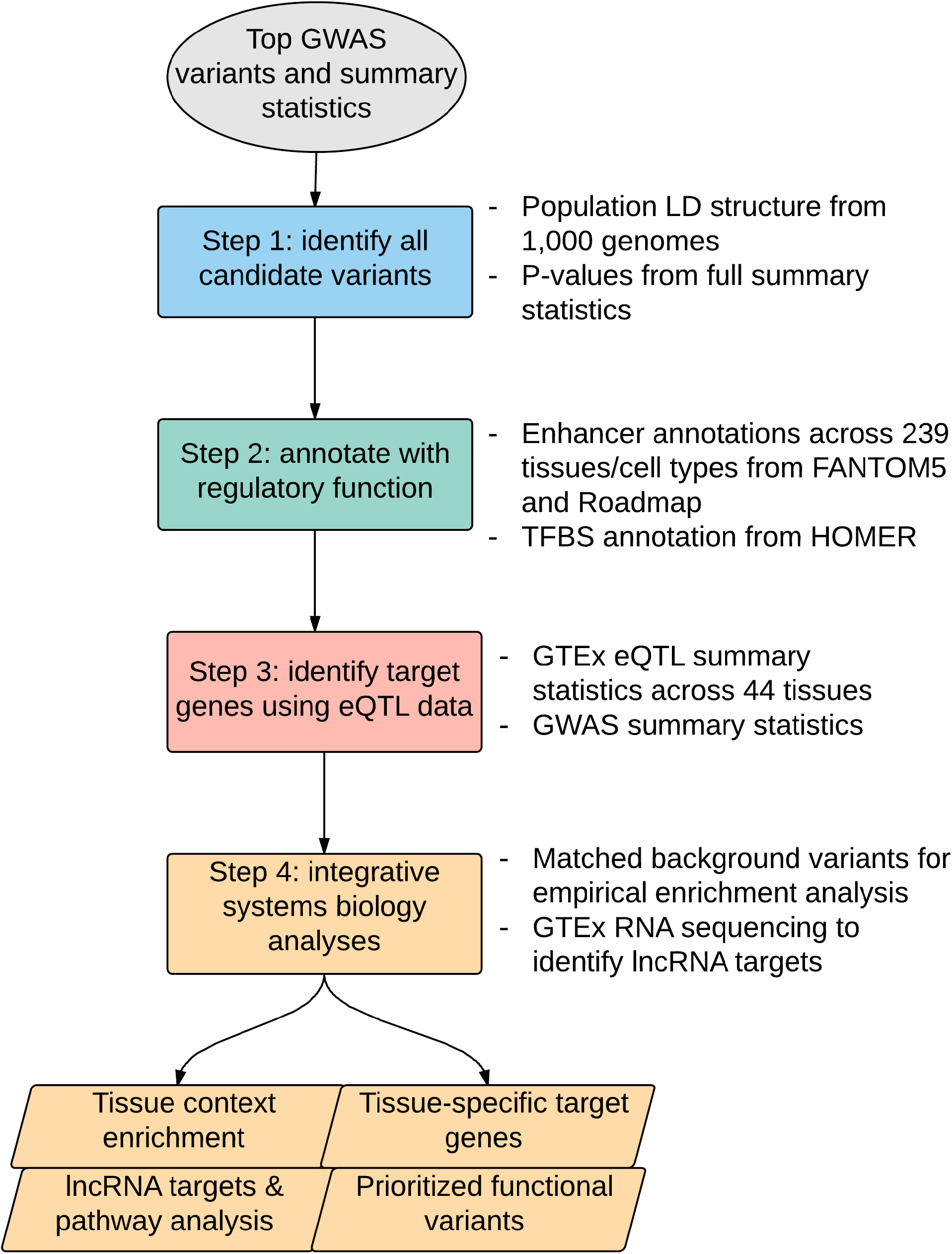
Outline of INFERNO pipeline approach.

### Defining comprehensive sets of potentially causal variants from GWAS findings

Given a user-defined list of top variants and summary statistics from a given GWAS study, INFERNO defines comprehensive sets of putatively causal variants underlying each top association signal (Methods). First, INFERNO uses GWAS summary statistics to identify all significant or almost significant variants near each tagging variant. This set is pruned by linkage disequilibrium (LD) structure using PLINK [18], where representative subsets of variants within each LD block are chosen such that subjecting them to LD expansion recaptures the other variants in the LD block. This analysis uses LD block data in a user-defined population (European, African, or Asian) from the 1,000 Genomes Project [19]. These LD block-tagging variants are re-expanded into full LD blocks, and these expanded sets are used for all further analysis to maximize the chance of finding the truly causal genetic variants underlying each association signal. This expansion approach provides a comprehensive set of putatively causal variants by taking full GWAS summary statistics as well as LD structure into account. If full summary statistics are not available, INFERNO can directly expand the top tag GWAS variants into LD blocks.

### Annotation of expanded variant sets with transcriptional regulatory elements

To identify noncoding genetic variants, INFERNO quantifies the proportion of variants overlapping messenger RNA (mRNA) promoters, exons (i.e. coding variants), introns, 5’ untranslated region (UTR) exons and introns, 3’ UTR exons and introns, and RepeatMasker genomic repeats [20] including LINE and SINE elements. Any variant outside all of these regions is classified as intergenic.

Next, each variant is overlapped with two complementary enhancer data sources. The first are sites of enhancer RNA (eRNA) transcription, which reflects enhancer activity [21], as assayed by cap analysis of gene expression (CAGE-seq) data generated by the FANTOM5 consortium across 112 tissue and cell type groupings [7]. The second is enhancer states defined by ChromHMM [22] using combinatorial epigenomic states measured by chromatin immunoprecipitation and sequencing (ChIP-seq) of 5 histone modifications, which mark active enhancers in a stereotypical pattern [23, 24], across 127 tissues and cell types generated by the Roadmap Epigenomics Project [8] and by the Encyclopedia of DNA Elements (ENCODE) project [9, 25]. In the Roadmap analysis, variants are overlapped with a total of 15 ChromHMM states including 3 types of enhancer states, sites of genic transcription, repressed regions, and active promoters, another type of transcriptional regulatory element that may harbor causal variants underlying an association signal.

In addition to overlapping variants with functional genomics annotations across tissues, INFERNO includes a sequence-based analysis to find variants affecting transcription factor binding sites (TFBSs) identified by the HOMER tool [26] (see Methods). INFERNO uses positional weight matrices (PWMs) to compute the difference in the log-odds binding probability of each affected TFBS between the reference and alternate alleles of the overlapping genetic variant (∆PWM score) in order to identify genetic variants that either increase or decrease TFBS strength.

### Tissue categorization of annotations and integrative analysis

Combining information across complementary sources of functional genomics data enables the comparison of evidence from independent experiments to improve sensitivity and robustness. However, it is not possible to directly compare results across the three consortia analyzed by INFERNO (FANTOM5, Roadmap, and GTEx) because each assayed different tissue types and cell lines at different levels of biological complexity. To integrate evidence from these disparate data sources, we designed a tissue classification scheme that grouped individual samples from each data source into one of 32 tissue categories (Supplementary Tables 1-3, Supplementary Figure 1). Datasets from both the GTEx and FANTOM5 consortia were grouped into high-level categories using the UBERON and CL ontologies for tissues and cell types, respectively [27, 28]. This integrative categorization provides a high-level view of the affected tissue contexts across consortia, allowing for easy identification of the tissue contexts harboring noncoding elements affected by variants within each GWAS tag region.

### Background sampling for enrichment of enhancer overlaps

Due to the widespread regulatory activity in the noncoding genome [9], selecting a large set of genetic variants in an LD block and overlapping them with hundreds of functional measurements may yield many overlaps simply by chance. To quantify the significance of enhancer overlap enrichment, INFERNO includes a statistical sampling approach to define empirical p-values for the enrichment of overlaps for each pair of annotations *a* and tissue category *t* within individual GWAS tag regions as well as across all tag regions (e.g. *a* = FANTOM5 enhancers, *t* = brain, see Methods).

### eQTL co-localization analysis

Current noncoding genetic variation annotation methods can identify functional variants and affected regulatory elements, and in some cases provide a hypothesis-free characterization of the relevant tissue context [29], but do not fully characterize the affected target genes. These methods either assume that the nearest gene is the affected transcript or directly overlap variants with eQTLs, including HaploReg, which directly overlaps variants with eQTL signals from 14 sources. However, the closest gene is typically not the target of transcriptional regulatory elements [5], and direct overlap of variants with eQTL signals is biased by genomic LD structures, where an eQTL association signal may be spread across a haplotype block so that the measured variant is not the causal regulatory variant.

When summary statistics are not available, INFERNO performs direct overlap with eQTL signals across 44 tissues from the Genotype-Tissue Expression (GTEx) project [15]. However, if summary statistics are available, INFERNO performs co-localization analysis using the COLOC method to control for bias from LD structures [16]. This model uses a Bayesian statistical model to calculate posterior probabilities for different causality relationships between GWAS and tissue-specific eQTL signals. The most relevant hypothesis from the COLOC output for INFERNO is H_4_, that there is a shared causal variant underlying both the eQTL signal and the GWAS disease signal. INFERNO performs co-localization analysis comparing all GWAS signals within 500kb of each tag variant with eQTL signals across all 44 GTEx tissues (Methods). Strongly co-localized signals are defined as those with P(H_4_) ≥ 0.5. COLOC also reports the probability of any individual variant being the shared causal variant, measured by the Approximate Bayes Factor (ABF). For further analysis of putatively causal variants, INFERNO defines sets of variants accounting for at least half of the cumulative ABF distribution at each strongly co-localized signal. This allows for the sensitive detection of truly co-localized signals to identify causal variants, the target genes they affect, and the tissue context of the regulation.

### Integrative analysis of co-localized eQTLs with annotations

To integrate the results between the enhancer and eQTL analyses, INFERNO uses the tissue categorizations of the FANTOM5, Roadmap, and GTEx datasets to stratify variants in the ABF-expanded sets underlying a co-localized signal by whether they affect a TFBS, overlap a FANTOM5 or Roadmap enhancer, and whether the enhancer came from the same tissue category as the eQTL signal. Variants in the ABF-expanded sets underlying a strongly co-localized eQTL signal overlapping enhancers from a concordant tissue class are prioritized if they also overlap a TFBS and/or have a high individual ABF value (Methods). This enables the identification of a small number of causal genetic signals and affected target genes supported by a variety of diverse functional genomics data sources all implicating the same tissue category. This integration of diverse data types spanning epigenomic marks, enhancer activity, transcription factor motifs, and eQTL signals provides a useful tool to identify causal variants, affected target genes, and relevant tissue contexts in an unbiased fashion. Furthermore, if users have an *a priori* assumption about which tissue categories might be relevant for their trait of interest, INFERNO can further prioritize variants affecting regulatory mechanisms from those specific categories. INFERNO provides several tables summarizing the annotation support for each co-localized signal as well as lists of gene symbols for usage in pathway analysis tools (Methods).

### Correlation-based lncRNA target identification

Finding a GTEx eQTL target gene supported by concordant enhancer support may be only the first step to understanding the affected regulatory mechanism underlying a genetic association signal, because the GTEx eQTL data also includes long noncoding RNA (lncRNA) signals, which can act as transcriptional regulatory elements for other genes [30]. Although tools for lncRNA target prediction exist, lncRNA targeting mechanisms are not fully characterized, so INFERNO takes an unbiased approach to finding their targets. RNA sequencing-based expression vectors of all genes in the genome including lncRNAs across all samples and tissues from GTEx are correlated with the expression vector of a lncRNA of interest. Then, genes with correlation values meeting user-specified threshold on Spearman and Pearson correlations (Methods) are considered to be putative lncRNA target genes, in line with previous approaches to lncRNA target identification [31]. This analysis is done automatically within INFERNO after the co-localization analysis is complete, and lists of lncRNA targets are provided for users to perform pathway analysis to characterize the affected biological processes downstream of the lncRNA signal, including lists of targets split by the GTEx tissue and tissue category of the lncRNA signal.

### Application to schizophrenia GWAS

To demonstrate INFERNO’s utility, we analyzed a GWAS dataset for schizophrenia from the Psychiatric Genomics Consortium with 108 LD-independent signals (n = 36,989 cases, 113,075 controls, [17]). P-value expansion of variants within an order of significance magnitude of the top variants yielded 3,778 unique variants, which were LD pruned down to 268 independent variants. LD re-expansion with a threshold of R^2^ >= 0.7 in the European population of 1,000 genomes then yielded 8,371 unique variants (Figure 2a). Genomic partition analysis of these variants supported their noncoding function, as only 79 (0.8%) of these variants were located in messenger RNA (mRNA) exons, with the majority in mRNA introns (3,050), repeat elements (2,819), or outside of any annotations (2,087) (Figure 2b). Overlapping these variants with enhancer annotations found widespread enhancer signals in the Roadmap data, with 4,127 (49%) variants overlapping a ChromHMM enhancer state in at least one tissue (Figure 2c). The FANTOM5 overlaps were limited due to the more conservative nature of the eRNA measurements, with 196 (2.3%) variants overlapping a FANTOM5 enhancer in at least one tissue. Finally, overlap with HOMER TFBSs found that 3,821 (45%) unique variants overlapped TFBSs for 233 unique transcription factors for a total of 10,798 variant—TFBS overlaps. The majority (8,976) of these overlaps lowered the predicted binding strength (Figure 2d). We observed no significant enrichments of enhancer overlaps in any tissue category when considering all the tag regions together. However, four individual tag regions (rs2239063 (12p13.33), rs2693698 (14q32.2), rs6704641 (2q33.1), and rs4330281 (3p24.3)) harbored significant enrichments of enhancers in categories related to immunity (blood, adipose cells, tonsils, immune organ) and stem cells (including iPSC and placenta) (Supplementary Figure 2). The enrichment in the immune system is of particular interest given the strong epidemiologic and molecular genetic evidence of immune dysfunction in schizophrenia [32, 33]. Next, 99,354 co-localization tests (COLOC) were performed across the 108 tag regions, identifying 969 unique tissue-target gene eQTL signals across all 44 GTEx tissues and 300 unique genes (including 57 lncRNAs from 34 tissues) that were strongly co-localized with schizophrenia GWAS signals (Supplementary Table 4, Supplementary Figure 3). To characterize the downstream biological processes affected by these variants, we performed pathway analysis on the 300 co-localized target genes using the WebGestalt tool [34, 35], but this did not yield any significant enrichments.

**Figure 2:**
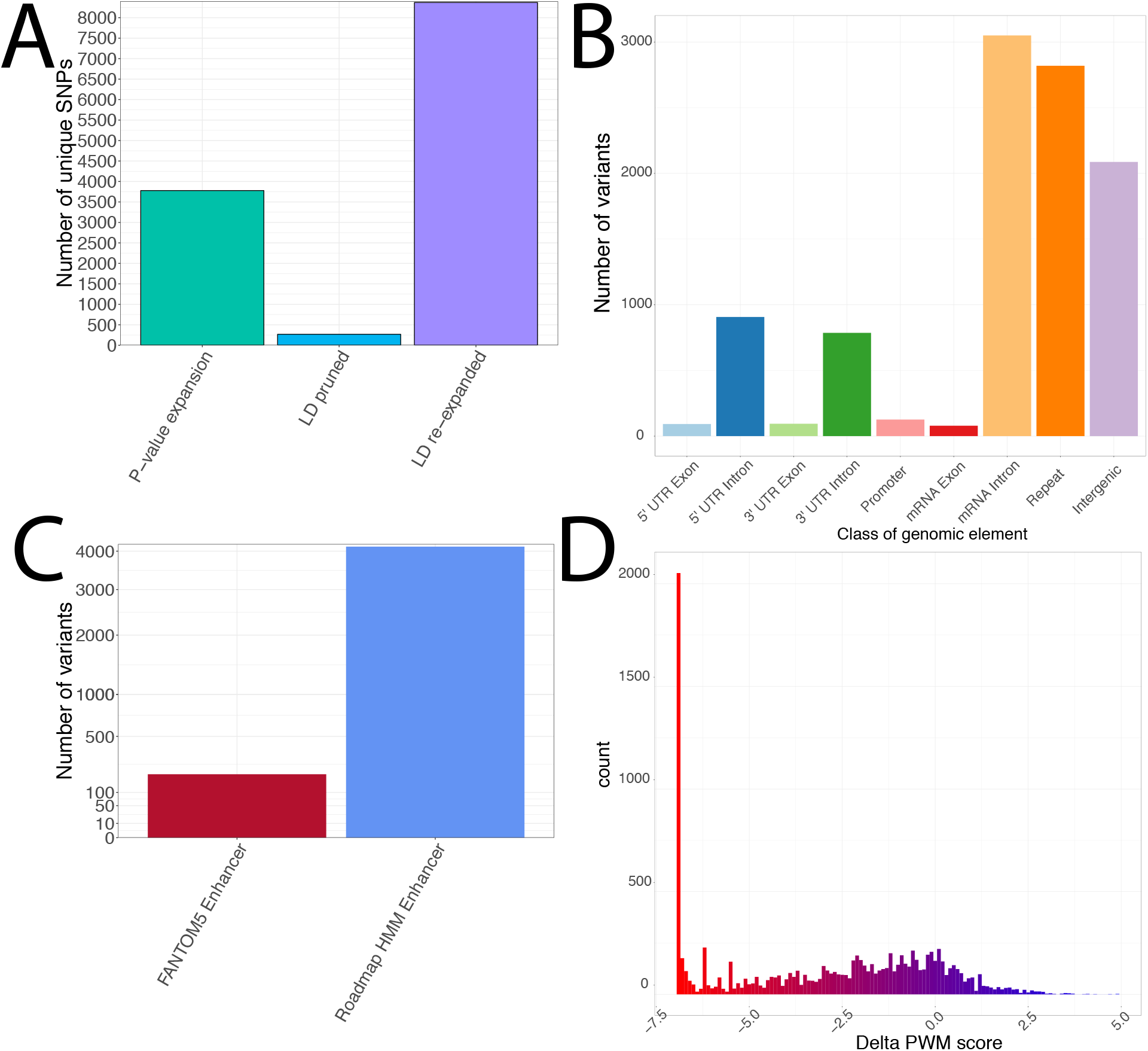
Characteristics of expanded variant sets for schizophrenia analysis. A) Number of variants after p-value expansion, LD pruning, and LD expansion. B) Counts of expanded set variants in genomic partitions. C) Number of variants overlapping enhancers from FANTOM5 and Roadmap. D) Distribution of ∆PWM scores for variants overlapping HOMER TFBSs.

Cross-reference of the significantly enriched tag regions with the COLOC results found that the rs2239063 region harbored co-localized eQTLs for *CACNA1C* in cerebellum and cerebellar hemisphere of brain (Supplementary Table 4). Variants in the ABF-expanded sets underlying these signals had concordant enhancer support from Roadmap enhancers in several data sources including thymus, hippocampus, and fetal brain as well as TFBS disruptions for the *RFX5* and *RFX1* transcription factors. These are two members out of 5 in the mammalian regulatory factor X (RFX) family of TFs, which play a key role in regulation of the immune response [36]. The brain microglial immune response plays a role in schizophrenia etiology and *CACNA1C* is a known schizophrenia-related gene [37, 38], supporting the utility and reproducibility of INFERNO results.

As an alternative approach for prioritizing the strongest results in the COLOC analysis, we identified 5 regions harboring co-localized eQTL and GWAS signals supported by variants with individually high ABFs as well as enhancer overlaps from the same tissue category: rs4766428 (12q24.11), rs4648845 (1p36.32), rs12826178/rs324017 (overlapping regions in 12q13.3), rs56205728 (15q15.1), and rs4702 (15q26.1) (Figure 3a, Supplementary Table 4). In the 12q24 region around rs4766428, rs4766428 itself was prioritized as having high ABF underlying 12 distinct eQTL signals including for *C12orf76* and *VPS29* in the brain category and *TCTN1* in the nerve tissue category (Figure 3b). Note that this variant lies in an intron of *ATP2A2* but does not target that gene. These three genes are all involved in transmembrane cellular processes: *C12orf76* is an unannotated transcript associated with the ‘ion channel activity’ GO pathway [39], *VPS29* is part of a group of vacuolar sorting proteins [40], and *TCTN1* encodes a family of secreted transmembrane proteins involved in ciliopathies and several cancer types [41]. This variant also disrupts binding sites for *ERRA, PPARg*, and *RXR* (∆PWM = -1.95, -1.84, -2.06, respectively). *ERRA*, also known as estrogen related receptor alpha, is an orphan nuclear receptor with no known ligand that is known to play a central role in regulating energy homeostasis and metabolism [42], which is disrupted in schizophrenia [43]. *PPARg* is a member of the peroxisome proliferator-activated receptor subfamily of transcription factors which are involved in adipocyte differentiation [44] in addition to having anti-inflammatory properties [45]. Furthermore, these factors form heterodimers with retinoid X receptors *(RXR)*, and the retinoid pathway has been suggested as a potential causal factor for schizophrenia [46, 47]. Of particular relevance is the ChromHMM enhancer overlap in brain dorsolateral prefrontal cortex (highlighted with red box in Figure 3b), the most relevant brain region in schizophrenia [48]. In the 1p36 region around rs4648845, a single variant, rs4592207, was found to underlie a pancreatic eQTL signal for the lncRNA *RP4-758J18.10* with high ABF. This variant also overlaps an *RFX5* binding site as well as one for *STAT3*, a member of the important signal transducer and activator of transcription (STAT) family of proteins. This protein family is associated with many human diseases, and drugs to inhibit *STAT3* phosphorylation are in clinical trials for the treatment of schizophrenia as well as several other diseases including Alzheimer’s disease, several cancer types, and type 2 diabetes [49].

**Figure 3:**
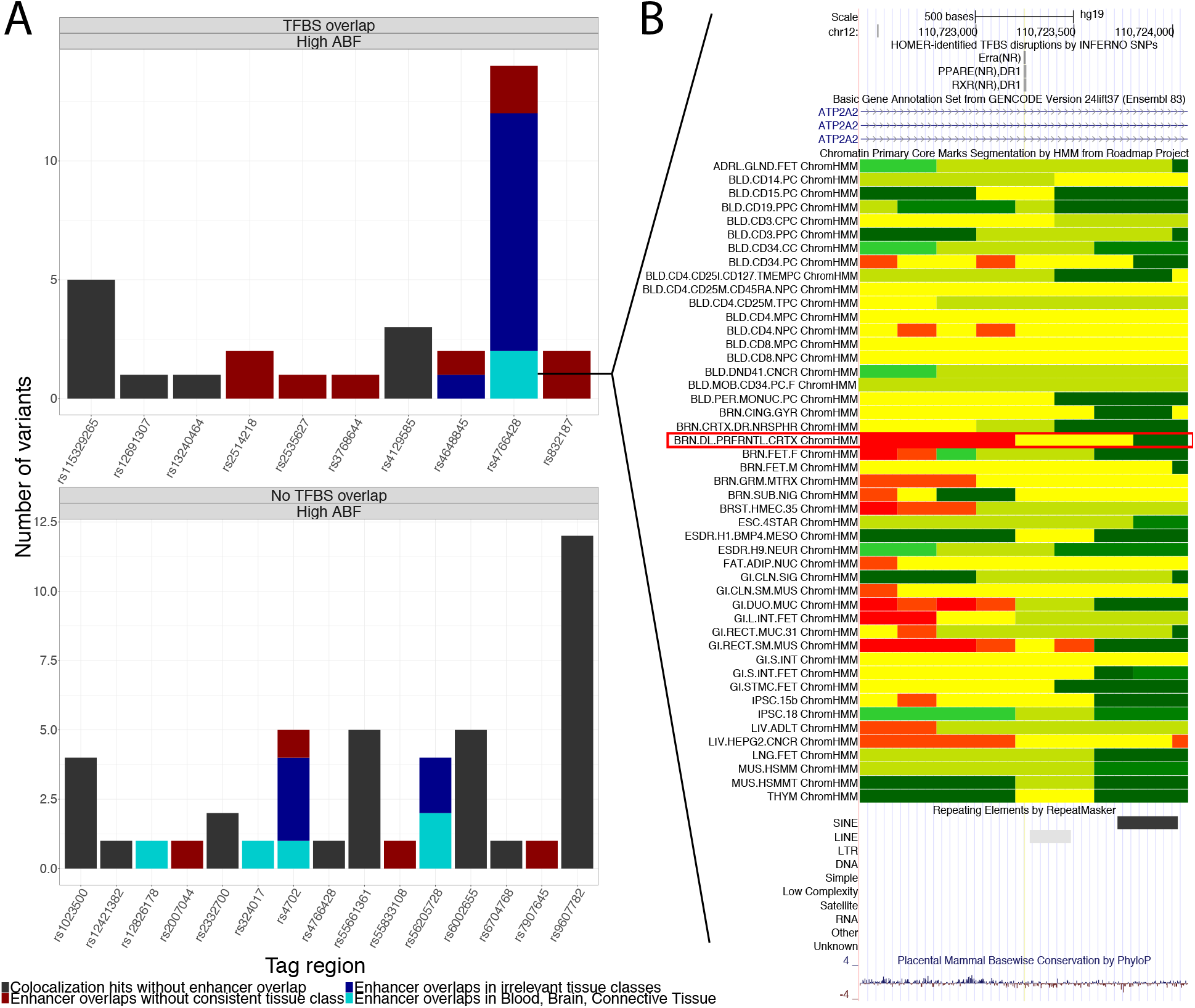
Results of GTEx co-localization analysis with schizophrenia GWAS. A) Top results from co-localization analysis integrated with annotation overlaps. Counts in barplots refer to individual variants underlying an eQTL signal in a given tag region, including all variants in the ABF-expanded sets. B) UCSC Genome Browser view of locus around rs4766428. In ChromHMM tracks, yellow = enhancer, green-yellow = genic enhancer, green = transcription, red = active transcription start site. Track highlighted with red box is dorsolateral prefrontal cortex.

In the 12q13 region around rs12826178, this analysis prioritized rs12826178 itself with high ABF underlying a blood eQTL signal for *TSPAN31*, another transmembrane protein also known as *SAS* that is amplified in human sarcomas [50]. This variant overlapped Roadmap enhancers from 9 tissues including several immune cells in the blood category but did not overlap any TFBSs. The variant rs12826178 was also prioritized in the nearby region around rs324017 for the same eQTL signal, but was not part of the LD expanded set in that region. In the 15q15 region around rs56205728, the variant itself was found to underlie 4 eQTL signals with high ABF, including *BUB1B* in brain cerebellar hemisphere and cerebellum and *PAK6* and *PLCB2* in skeletal muscle. This variant overlaps Roadmap enhancers from 28 data sources including astrocytes and several immune cell lines. *BUB1B* is a mitotic checkpoint serine/threonine kinase, and is differentially overexpressed in both schizophrenia and bipolar disorder brains [51]. *PAK6* is another serine/threonine kinase that is a member of the p21-activated kinase (PAK) family. PAK inhibitors ameliorate dendritic spine deterioration associated with schizophrenia [52], and disruption of this gene in mice leads to learning and locomotion deficits [53]. *PLCB2* is a phosphodiesterase that is involved in innate immunity [54]. Finally, in the 15q26 region around rs4702, rs4702 is the prioritized variant underlying four eQTL signals, for *FES* in subcutaneous adipose and pancreas, *SLCO3A1* in transformed fibroblasts, and *FURIN* in esophagus mucosa. This variant overlaps enhancers from 46 Roadmap datasets, and although the tissues do not seem to be directly relevant to schizophrenia, there is experimental evidence that *FURIN* is involved in neurodevelopmental processes that may be affected in schizophrenia [55].

Using the top 108 LD-independent signals as input, HaploReg detected almost none of the top results identified by INFERNO. In the rs2239063 region, no eQTLs for *CACNA1C* were detected. In the rs4766428 region, HaploReg did not identify any brain signals for *VPS29* and did not identify any eQTL signals at all for *C12orf76* and *TCTN1.* In the rs4648845 region, the variant we prioritized, rs4592207, is not part of the LD-expanded block analyzed by HaploReg. In the rs1286178 region, HaploReg did not detect any eQTL signal for rs1286178. In the rs56205728 region, HaploReg did not identify the brain signals for *BUB1B* or the *PAK6* signal, although it did detect a *PLCB2* eQTL, albeit in lymphocytes and lung rather than the skeletal muscle signal INFERNO prioritized. Finally, in the rs4702 region, HaploReg detected the *FES* signals in pancreas but not subcutaneous adipose, detected the *FURIN* signal in esophagus mucosa, and missed the *SLCO3A1* fibroblast signal, although it identified additional *FES* signals in fibroblasts and thyroid that INFERNO did not identify as strongly co-localized signals (P(H_4_) = 0.08 and 0.40, respectively).

Next, we performed correlation-based target identification for the lncRNAs targeted by co-localized eQTL signals, which identified 6,005 unique genes targeted by 46 unique lncRNAs from 33 tissues and 15 tissue classes (Supplementary Figure 4). We first performed pathway analysis on all 6,005 genes targeted by these lncRNAs. This found significant enrichments in several schizophrenia-related pathways (Supplementary Table 5) including RNA splicing [56], phosphatidylinositol signaling [57], Th1 and Th2 cell differentiation [58], T cell receptor signaling [59], and RNA transport [60].

To refine our understanding of the downstream effects of these lncRNAs, we split the lists of target genes by which tissue category the lncRNA eQTL signal came from and performed pathway analysis separately in each category (Figure 4). This identified tissue-specific pathway effects in pathways known to be related to schizophrenia such as the *MAPK* signaling pathway (KEGG pathway ID hsa04010) [61] in the blood category as well as pathways that were enriched across several contexts, notably implicating the spliceosome (hsa03040) [56], which was enriched in 10 categories including blood and brain. Another intriguing signal was the enrichment of the Herpes simplex infection (hsa05168) pathway in 8 categories, also including blood and brain. Maternal Herpes simplex virus (HSV) infection may lead to increased risk of schizophrenia in their offspring [62] and HSV exposure may exacerbate cognitive function impairment in schizophrenic patients [63]. Thus, the lncRNA target identification performed by INFERNO can identify biologically relevant genes and pathways downstream of lncRNA perturbations by genetic variants.

**Figure 4:**
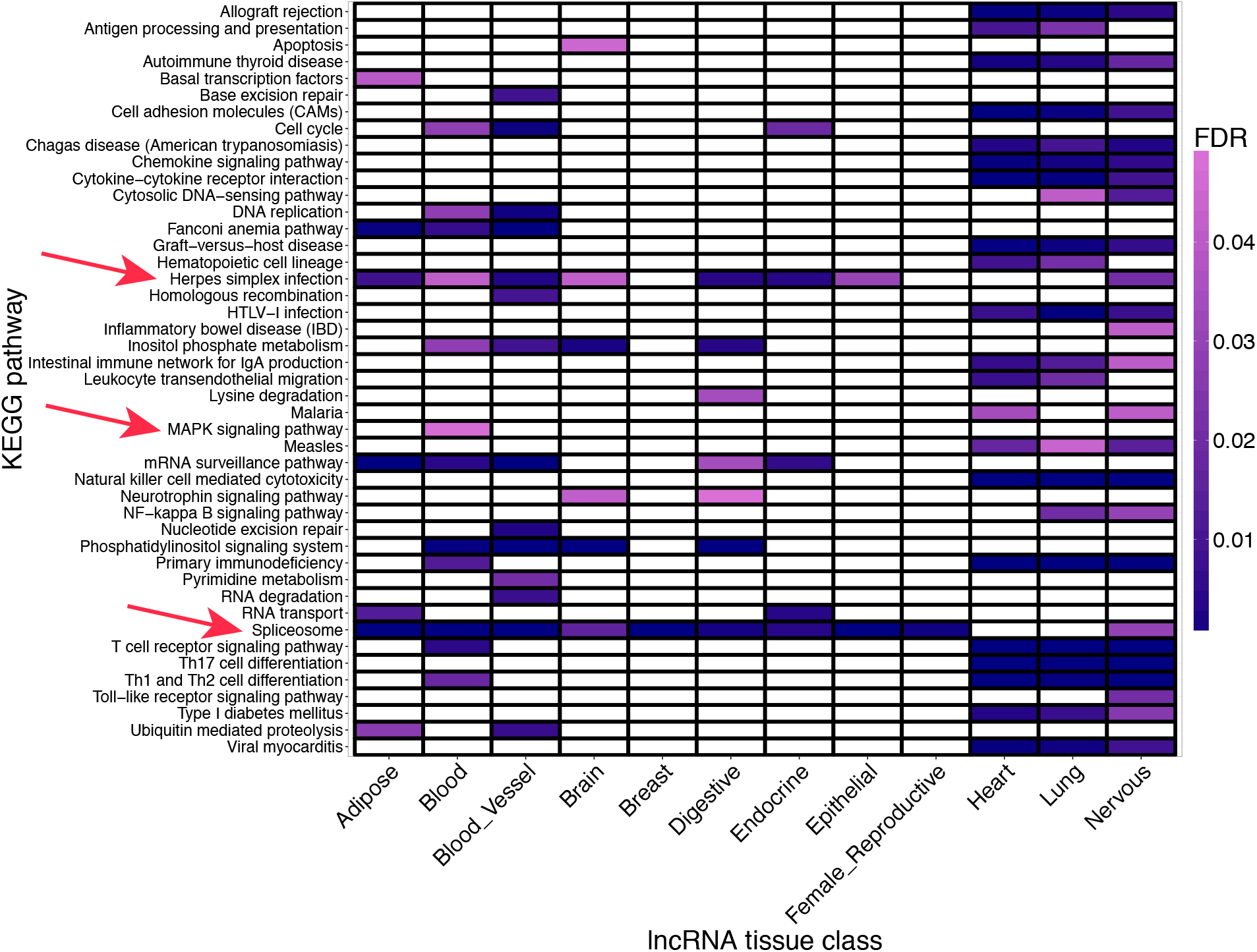
KEGG pathway enrichments for lncRNA targets. Results are split by the tissue category of the lncRNA eQTL signal. Red arrows denote known schizophrenia-related pathways discussed in the main text.

### Web server and tool availability

We provide a web server for INFERNO analyses that accepts the top variants from any given GWAS and performs the annotation overlap analysis including directly overlapping variants with GTEx eQTL data [64]. To run the computationally intensive enhancer sampling, eQTL co-localization, and lncRNA correlation analyses, INFERNO is also available as an open source pipeline [65].

## Discussion

INFERNO provides a sensitive hypothesis generation method for identifying functional genetic variants underlying genetic association signals and characterizing their tissue-specific effects on regulatory elements, target genes, and downstream biological processes. The schizophrenia analysis demonstrated that INFERNO picks up many signals that converge to common tissue contexts and pathways when sufficient genetic loci are available. However, while the diversity of functional genomic data and tissue contexts analyzed by INFERNO allows it to identify functional variants, regulatory elements, and target genes underlying GWAS association signals, this broad range of data sources also means that our algorithm may pick up more general regulatory mechanisms not directly related to the phenotype of interest, and these “hitchhikers” could obfuscate the truly causal processes. This is likely to be a characteristic of complex traits in general. Another factor that affects the specificity of INFERNO results is the currently limited availability of functional genomics annotation data, which are measured in normal tissues that do not reflect the disease state for a given GWAS signal and may not be exact matches for the relevant tissue context for a given trait. Thus, INFERNO is best used as a powerful tool to prioritize biological processes and tissue contexts in an unbiased and systematic fashion for functional follow-up studies to prove the causality of the prioritized signals and their relevance to the phenotype of interest.

INFERNO improves on existing noncoding annotation methods for GWAS signals, the most comparable of which is HaploReg [14]. HaploReg expands GWAS variants by LD structure only, missing many of the candidate variants INFERNO identifies using summary statistic-based expansion, and reports direct annotation overlaps with Roadmap but not FANTOM5 enhancer annotations. Additionally, it lacks a tissue classification framework to integrate information across disparate annotation sources. HaploReg provides an enhancer enrichment score by calculating the background frequencies of enhancer overlap in each cell type for all unique GWAS loci and all 1,000 Genomes common variants and comparing these frequencies to those for a query list of variants using a binomial test. This approach ignores LD structure and does not match variants by any characteristics. INFERNO provides a more sensitive statistical method for quantifying the tissue-specific significance of annotation overlaps in a GWAS signal accounting for LD structure and other genomic characteristics. Furthermore, INFERNO allows for the calculation of enrichments both within and across tag regions, and the tissue classification approach enables the scoring of enrichments supported by disparate data sources. INFERNO also performs a more sensitive eQTL analysis by applying a Bayesian model to identify truly co-localized signals between GWAS and eQTL data, and additionally performs target identification for lncRNA targets identified by this algorithm.

Application of INFERNO to schizophrenia GWAS data identified significant overlaps of enhancers in immune- and brain-related tissue categories and eQTL signals from the same categories targeting known schizophrenia genes. This analysis identified putatively functional variants in 6 tag regions and also identified tissue-specific lncRNA signals targeting several biological processes known to be related to schizophrenia including the *MAPK* signaling pathway, spliceosome, and Herpes simplex infection.

INFERNO is limited by currently available functional genomics datasets, and will continue to increase in power as functional datasets are generated in more tissue and cell types and as genomics technology continues to be refined and improved. Another movement in the field is to consider structural variation and copy number variants in addition to the single nucleotide variants that INFERNO currently analyzes, and as more of these data are generated INFERNO will be updated to allow for the analysis of these signals in the noncoding genome.

## Conclusion

In this manuscript, we present the INFERNO (INFERring the molecular mechanisms of NOncoding genetic variants) tool for characterizing functional variants underlying GWAS association signals as well as the regulatory mechanisms, tissue contexts, target genes, and biological processes they affect. This tool integrates information across hundreds of functional genomics datasets to provide a data-driven, hypothesis-free approach for detailed characterization and identification of noncoding genetic variants underlying genetic association signals to provide biologically interpretable results. This characterization of the relevant tissue contexts and biological pathways underlying disease risk can improve the mechanistic understanding of disease risk as well as identify candidate therapeutic targets for future pharmacological interventions.

INFERNO provides the most comprehensive and unbiased tool to identify causal noncoding variants disrupting enhancers and the downstream effects of these disruptions including the relevant tissue context and affected target genes and pathways. We provide a web server (http://inferno.lisanwanglab.org) that takes in top GWAS variants, expands them into LD blocks, and annotates them with functional genomics data and direct eQTL overlap. We also provide open source code [65] for the full pipeline that runs on full GWAS summary statistics and performs p-value as well as LD expansion, sampling for functional enrichment, GTEx eQTL co-localization, and lncRNA target identification in addition to the annotation overlaps provided by the web server.

## Methods

### P-value and LD expansion of IGAP variants

Given GWAS summary statistics and a set of user-defined top variants, INFERNO first computes the sets of all variants *i* within 500 kb of each tagging variant such that *p*_*t*_ ≤ *m * p*_*t*_ where *p*_*i*_ is the p-value assigned to variant *i, p*_*t*_ is the p-value of the tagging variant, and *m* is the user-defined multiplicative constant, 10 by default for one order of magnitude. These sets are pruned by linkage disequilibrium using PLINK v1.90b2i 64-bit [18] with the parameters --vcf for the input file and ‘--indep-pairwise 500kb 1 0.7’ (within 500,000bp and meeting a correlation threshold of R^2^>= 0.7). LD structure information is calculated using the phase 3 version 1 (May 11, 2011) of the 1,000 Genomes Project [19]. For the schizophrenia analyses in this manuscript, data from the European (EUR) population was used, but 1,000 Genomes also provides LD structure information for Africa (AFR), Asia (ASN), and the Americas (AMR) and INFERNO allows users to choose their population of interest. Then, variants are re-expanded by LD structure using the parameters --vcf for the file containing neighboring variants and ‘--allow-no-sex, --r2 with-freqs dprime --ld-snp RSID --ld-window 99999 --ld-window-kb AREA -ld-window-r2 R2’, where RSID is the tag variant of interest, AREA is the LD block size threshold parameter, and R2 is the parameter defining the threshold on R^2^.

### Genomic partition analysis

Variants were categorized into different functional categories using the UCSC knownGene and RepeatMasker annotations for the hg19 genome build. Only chr1-22, X and Y are used in INFERNO. The 5’ UTR exons and introns, 3’ UTR exons and introns, and exons and introns were extracted from the knownGene annotation for each protein-coding gene, and all overlapping exons were merged together. Promoter annotations were defined as 1,000bp upstream of the first exon in the transcript, either coding or in the UTR. Variants were then assigned to mutually exclusive genomic element annotations using the hierarchy: 5’ UTR exon > 5’ UTR intron > 3’ UTR exon > 3’ UTR intron > promoter > mRNA exon > mRNA intron > repeat. A variant not overlapping with any class of elements above was classified as intergenic.

### Functional annotation data download and pre-processing

FANTOM5 enhancer facet-level expression BED files, Roadmap ChromHMM BED files for the 5 core Roadmap marks (H3K4me3, H3K4me1, H3K36me3, H3K27me3, H3K9me3), HOMER TFBS annotations, and GTEx eQTL and RNAseq data were downloaded from their respective servers and further processed using the bedtools suite of tools [66] and custom awk and Python scripts. A detailed description of the data sources and pre-processing steps is available at http://inferno.lisanwanglab.org/README.html and the full processed annotation data is available in the ‘Availability of Data and Materials’ section.

### INFERNO pipeline implementation

The INFERNO pipeline is implemented using Python v2.7.9, R v3.2.3, and bash, and is available at https://bitbucket.org/alexamlie/inferno. The pipeline allows user-defined parameters for the 1,000 Genomes population to use, the order of magnitude cutoff for p-value expansion, the LD and distance thresholds to use for LD expansion, the size of the window around FANTOM5 loci, and whether or not to calculate ∆PWM scores. It is also flexible and can run any or all of the analysis steps including p-value and LD expansion, genomic partition analysis, overlap with FANTOM5, direct overlap with GTEx eQTLs, overlap and calculation of ∆PWM scores with HOMER-defined TFBSs, annotation with ChromHMM states across all 127 Roadmap tissues and cell types, statistical analysis of enhancer enrichment, eQTL co-localization, and lncRNA target identification.

### Dataset classification into tissue categories

Building off the existing categorization of Roadmap samples and informed by the UBERON and CL ontologies [27, 28] used in the FANTOM5 facet-level classification and in GTEx, the different tissues and cell types from each data source were grouped into 32 major classes used for this analysis, and some of the data sources were further grouped into 58 secondary and 15 tertiary sub-classes (Supplementary Table 1).

### Quantification of enhancer enrichments

10,000 random sets of background variants matched to the input set of variants (before LD expansion) by distance to the nearest gene, minor allele frequency, and the number of variants in each tagged LD block are sampled. The variants from each background set are then expanded into their corresponding LD blocks to match the number of variants in the LD expanded input set and overlapped with the same sets of functional annotations. Then, the empirical *p*-value for the significance of the overlap of the input data with each functional annotation or combination of annotations *a* (e.g. eRNA enhancer overlap, or both eRNA enhancer overlap and eQTL overlap) in a given tissue category *t* is defined as 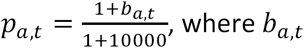 is the number of background samples that include at least as many LD blocks overlapping the annotation *a* in the tissue context *t* as the input dataset. By default, each LD block only contributes one effective count for annotation overlap in order to correct for LD structure, but INFERNO will also report results counting each variant in an LD block separately. These empirical p-values are corrected for multiple testing using the Benjamini-Hochberg procedure [67].

### eQTL Colocalization analysis

For colocalization analysis, INFERNO uses the COLOC R package [16] to compare the eQTL signals tested in GTEx across all 44 tissues against GWAS summary statistics. For each tag region and GTEx tissue, the script identifies all the genes tested for eQTL with the tagging variant in the region, reads in the eQTL data for each gene, and performs colocalization analysis using all the GWAS variants 500,000bp on either side of the tag variant that are also found in the eQTL data. Minor allele frequencies (MAFs) can be defined by the user or can be extracted from 1,000 Genomes data using a custom preprocessing script. Then, the MAF and p-values of variants in the GWAS and eQTL datasets are used for co-localization analysis, including a user-defined sample size and case/control ratio for the GWAS of interest.

### lncRNA correlation analysis

To find the target genes of each lncRNA found to have a co-localized eQTL signal, reads per kilobase per million (RPKM) values across all RNA-sequenced samples in GTEx are used. GENCODE annotations are used to identify GTEx target genes that are categorized as lncRNAs [68], and the RPKM expression files are used to extract the expression of all genes expressed in at least one sample, including each lncRNA, for one chromosome at a time. Then the pipeline calculates the correlations of the lncRNA expression vectors against all columns (genes) of the gene expression matrix using the corr.test function from the psych R package [69]. Two correlation measures are computed: the Pearson correlation, which measures the linear relationship between two variables, and the Spearman correlation, which is a rank-based test that does not assume a linear relationship. User-defined parameters on the absolute value of both the Spearman and Pearson correlation measures, 0.5 by default, are used to identify lncRNA target genes.

### Schizophrenia GWAS analysis

The full summary statistics file (scz2.snp.results.txt) and 128 top variants (scz2.rep.128.txt) for the schizophrenia analysis were obtained from the Psychiatric Genomics Consortium downloads page [70]. The top variants were parsed to remove variants on sex chromosomes and converted into INFERNO input format using awk scripts. The summary statistics were annotated with minor allele frequencies from the 1,000 genomes data using a custom script, annotate_input_variants.R, in the data_preprocessing/ section of the INFERNO code. These parsed files were then used as input to INFERNO, and the exact call used is available in the INFERNO README file [65].

## Declarations

### Ethics approval and consent to participate

Not applicable.

### Consent for publication

Not applicable.

### Availability of data and materials

The datasets supporting the conclusions of this article are available in the full INFERNO annotation file available at http://tesla.pcbi.upenn.edu/~alexaml/INFERNO/full INFERNO annotations.tar.gz. The web server is available at http://inferno.lisanwanglab.org/. The INFERNO software is open source and available at https://bitbucket.org/alexamlie/INFERNO/. The full INFERNO pipeline runs on Unix, is implemented using bash, Python 2.7, and R, and runs on bsub-based cluster computing systems. Further software versions, specific annotation sources and pre-processing scripts, and package requirements are documented in the Bitbucket repository and web server. INFERNO is licensed under the MIT license.

## Authors’ contributions

AAW, MT, and LSW initially conceived of the overall INFERNO pipeline approach. AAW implemented the INFERNO pipeline and oversaw the majority of the development, implemented the web server, analyzed the schizophrenia data, and wrote the manuscript. EEM performed exploratory network analysis and contributed to interpretation of the schizophrenia results. PPK and OV helped set up the web server and provided input on the user interface. ZK helped with the graphic design of the web server. DT provided expert advice and interpretation for the schizophrenia results. CDB, GDS, and LSW provided valuable advisory input throughout the development process and helped to acquire data to test INFERNO with. All authors read and approved the final manuscript.

## Acknowledgements

We gratefully acknowledge Chris Stoeckert for guidance on the tissue categorization, and all members of the Wang lab for valuable feedback, especially Fanny Leung.

---

**Supplementary Figure 1:**
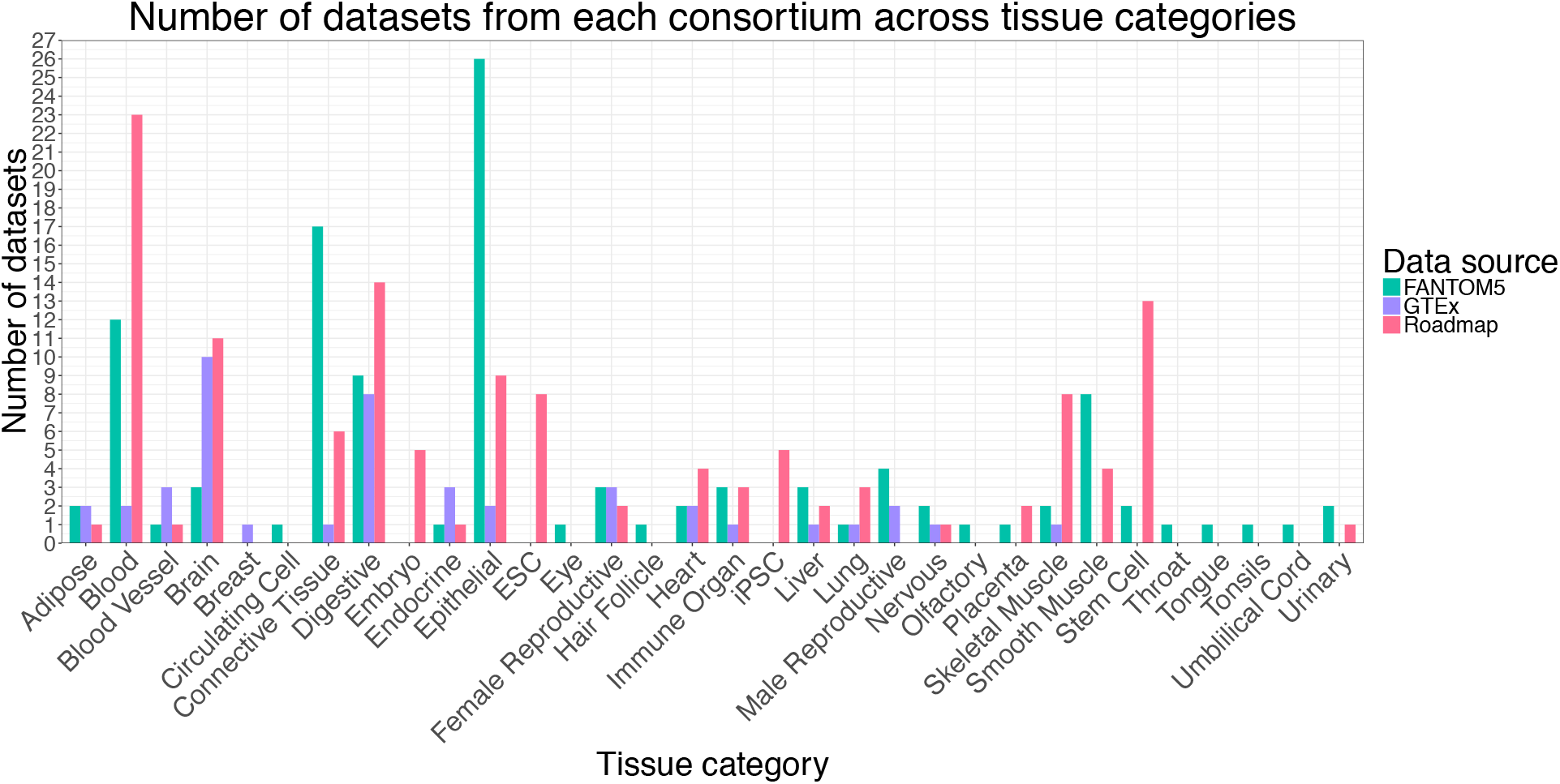
Barplot of counts of individual data sources grouped into each tissue category by consortium.

**Supplementary Figure 2:**
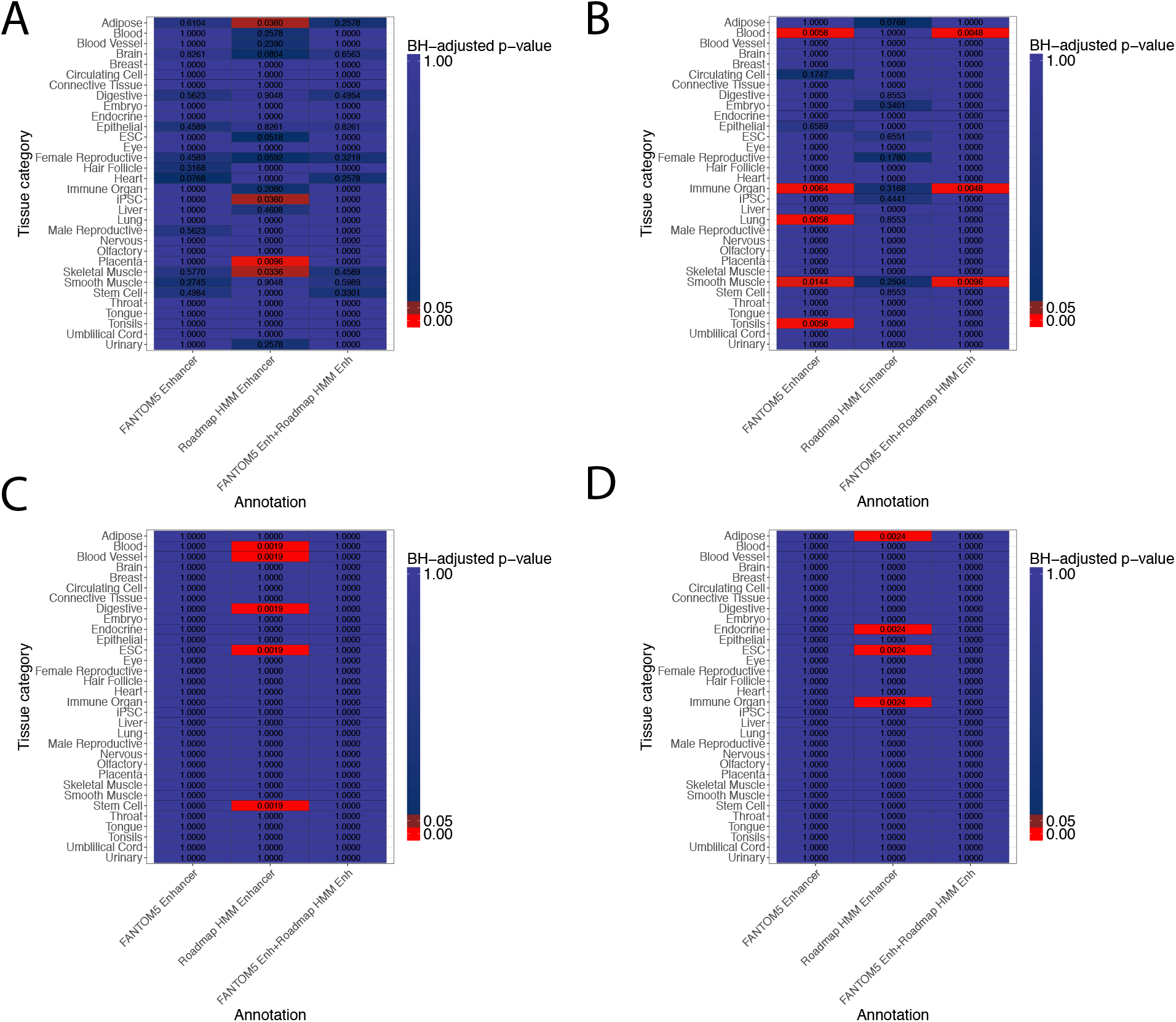
Tag region-specific sampling enrichment results for schizophrenia signals. A) Region around rs2239063 (12p13.33). B) Region around rs2693698 (14q32.2). C) Region around rs6704641 (2q33.1). D) Region around rs4330281 (3p24.3)

**Supplementary Figure 3:**
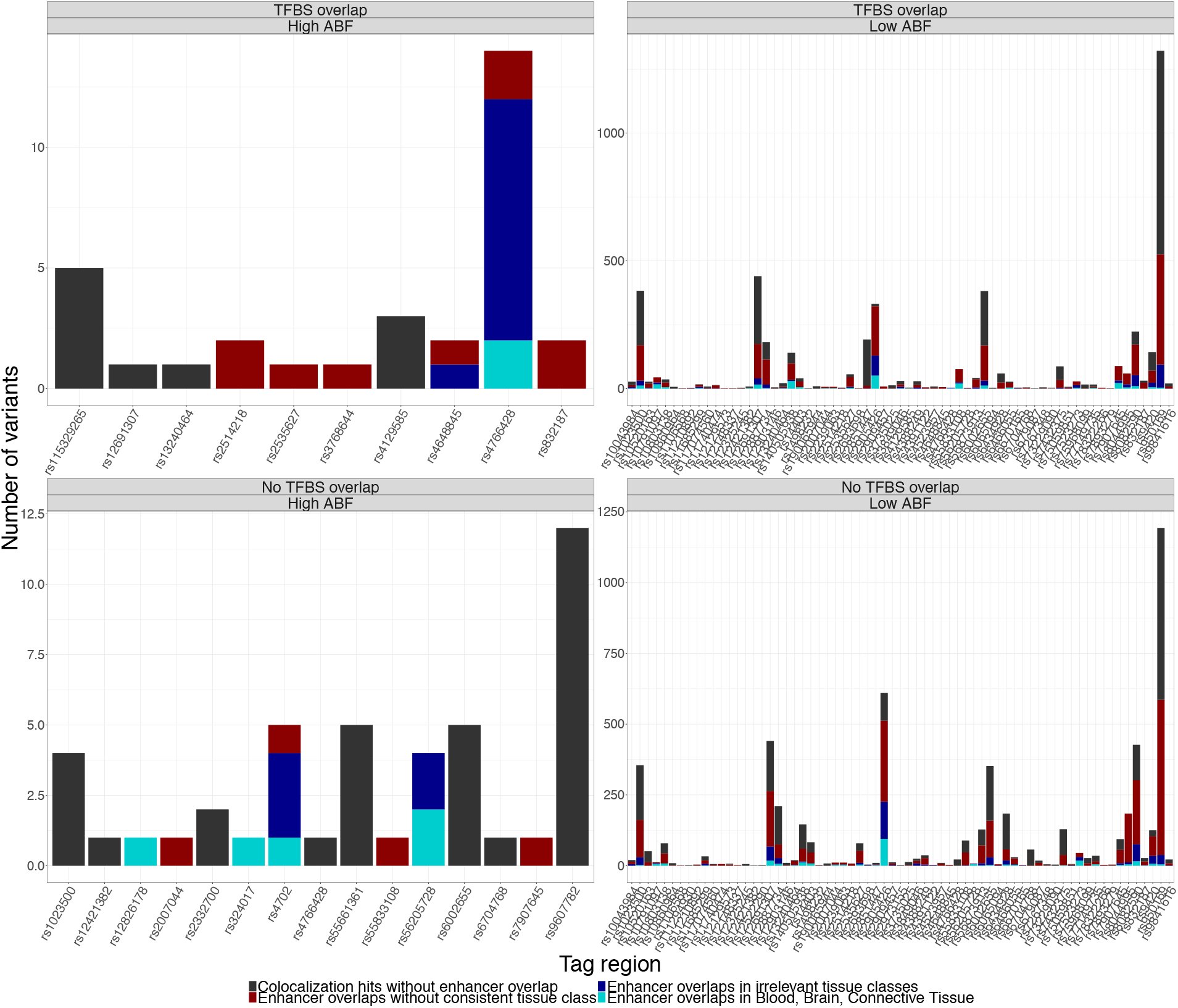
All results from co-localization analysis integrated with annotation overlaps. Counts in barplots refer to individual variants underlying an eQTL signal in a given tag region, including all variants in the ABF-expanded sets.

**Supplementary Figure 4:**
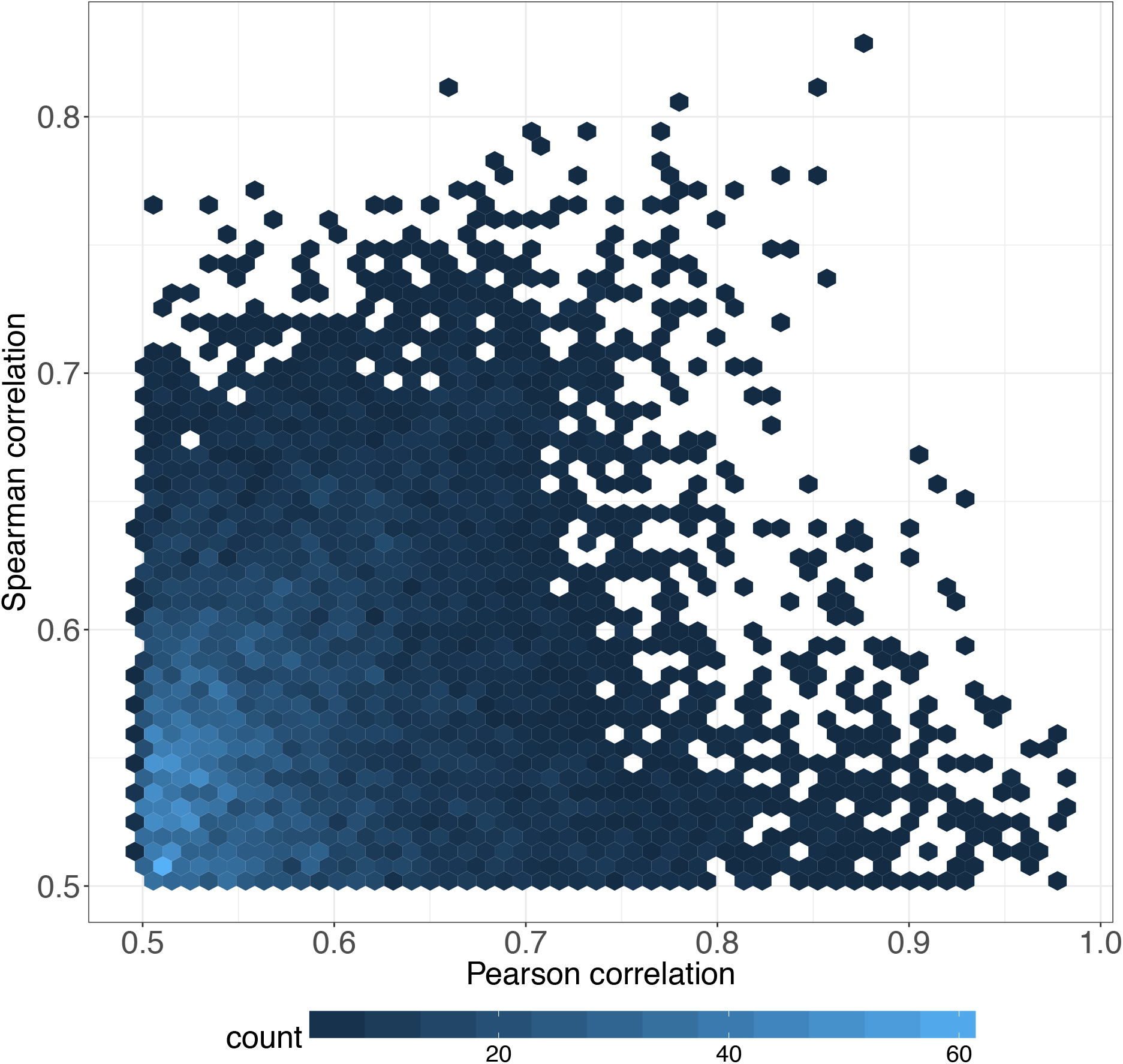
Scatterplot of distribution of Pearson vs. Spearman correlations of top mRNA transcripts correlated with lncRNAs identified as eQTL targets by INFERNO.

